# Asymmetric phosphoinositide lipid bilayers generated by spontaneous lipid insertion

**DOI:** 10.1101/2025.03.26.645509

**Authors:** Gwendal Guerin, Thi Lan Anh Nguyen, John Manzi, Christophe Le Clainche, Julien Heuvingh, Julien Berro, Nicolas Rodriguez, Sophie Cribier, Feng-Ching Tsai

## Abstract

Phosphatidylinositol phosphate (PIP) lipids, enriched on the cytoplasmic leaflet of the plasma membrane, are key regulators of diverse cellular processes, often through interactions with partner proteins that regulate actin assembly. Supported lipid bilayers (SLBs) provide a powerful model system to study the interactions of PIP lipids with their partner proteins. However, despite advances in SLB preparation methods, it remains a challenge to robustly obtain fluid SLBs in which PIP lipids are both mobile and asymmetrically distributed. In this study, we report a simple and robust method to generate asymmetric SLBs containing tunable amounts of PI(4,5)P_2_. By dissolving PI(4,5)P_2_ below its critical micelle concentration (CMC), we enable its spontaneous insertion into the upper leaflet of SLBs exposed to bulk solution. The mobility of PI(4,5)P_2_ is confirmed using fluorescence recovery after photobleaching (FRAP). Furthermore, we demonstrate that PI(4,5)P_2_ incorporated using this method retains its functionality, recruiting binding partners, actin-membrane linker ezrin, and myosin 1 motors capable of sliding actin filaments on the SLBs. Our method offers a straightforward strategy to generate asymmetric PI(4,5)P_2_-containing SLBs and is applicable to other lipid species with high CMC values.

## Introduction

The asymmetric distribution of lipid species is a fundamental feature of the plasma membrane in eukaryotic cells^1^. Sphingomyelin (SM) is abundant in the exoplasmic leaflet, whereas phosphatidylethanolamine (PE), phosphatidylserine (PS), and notably phosphatidylinositol (PI) lipids are predominantly found in the cytoplasmic leaflet^1^. PI lipids account for ∼10% of total lipids in mammalian cells^2^. Despite being a minor lipid species in the plasma membrane, PI and its phosphorylated derivatives, phosphatidylinositol phosphate (PIP) lipids, are recognized as key cellular regulators, affecting processes such as membrane trafficking and actin cytoskeleton assembly^3,4^. At neutral pH, PIP lipids exhibit a greater negative charge at the head group than PS, due to the presence of two negative charges per phosphate^5^. The phosphate modification enables PIP lipids to interact specifically with phosphoinositide-binding protein domains including pleckstrin homology (PH), Phox (PX), and BAR domains^6–8^. Among the PIP lipids, PI(4,5)P_2_ (hereafter referred to as PIP_2_) is the most abundant, accounting for 0.5 up to 4 mol% of the total plasma membrane lipids, estimated to be ∼ 4000 up to 34, 000 molecules per mm^2^ ^5,6^. By interacting with hundreds of effector proteins, including actin-binding proteins, PIP_2_ plays a central role in regulating numerous membrane-associated cellular processes^4,5,9,10^. PIP_2_ has therefore been extensively studied, especially through in vitro reconstitution assays using model membranes. For instance, PIP_2_ has been shown to form clusters by interacting with positively charged divalent ions such as Ca^2+^ and Mg^2+^ ^11–14^. Moreover, interactions between PIP_2_ and effector proteins, including members from the ezrin-radixin-moesin (ERM) protein family and actin nucleation promoting factors, have been implicated in the regulation of actin assembly^4,9,10,15–19^. However, despite these significant findings, how the dynamic interactions between PIP lipids and their effector proteins give rise to the diverse membrane processes in cells remains to be fully resolved.

Supported lipid bilayers (SLBs) are widely used model membrane platforms for studying lipid-lipid and protein-lipid interactions. Thanks to the substrate support and the planar shape, SLBs are particularly well suited for experimental techniques that require the imaging plane to be close to the substrate, such as atomic force microscopy, total internal reflection fluorescence (TIRF) microscopy, fluorescence correlation spectroscopy (FCS), single molecule tracking, and super-resolution imaging^20–24^. Common methods for generating SLBs include Langmuir-Blodgett deposition, the fusion of small vesicles (of the order of 100 nm in diameter), the rupture of giant vesicles (of the order of 10 μm in diameter), and rapid solvent exchange to drive SLB formation^25–28^. Vesicle fusion and rupture can produce symmetric SLBs (sym-SLBs), in which the lipid composition is the same on both leaflets. However, a key limitation with sym-SLBs is the inherent and often undesirable interactions between the lipids and the substrates. These lipid-substate interactions are known to reduce lipid mobility, likely due to factors such as surface roughness and electrostatic interactions^29–32^. Such interactions are particularly pronounced for negatively charged lipids like PIP_2_, which can become nearly immobilized in the bottom lipid leaflet that contacts the glass substrate^31,33–36^. Such immobilization compromises the quantitative analysis of lipid dynamics, and can also affect lipid lateral organization, such as phase separation, in the bilayer^37–39^.

To mitigate substrate-induced lipid immobilization, one approach is to replace glass substrates with mica, which provides a smoother and more hydrophilic surface compared to glass^29^. Alternatively, introducing polymer cushions, such as polyethylene glycol (PEG), between the substrate and bilayer can reduce friction^40,41^. Moreover, suspended planar lipid bilayers generated in micropores are elegant approaches that avoids the use of solid substrates^42–45^. However, these approaches often involve complex surface modifications or microfabrication^41–45^.

To enable lipid-protein interactions while minimizing substrate-induced immobilization, an effective approach is to construct asymmetric SLBs (asym-SLBs) in which the lipid species of interest, such as PIP_2_, are enriched in the top leaflet facing the bulk solution. Langmuir-Blodgett deposition can be used to generate asym-SLBs, but it requires sophisticated procedures^26^. Other methods include methyl-β-cyclodextrin-assisted lipid exchange to enrich sphingomyelin in the top lipid leaflet facing to the buffer^46,47^, using negatively charged surfaces to promote a higher amount of negatively charged lipids in the top leaflet^48^, transferring lipids to the top leaflet by the hemifusion of vesicles on the SLBs^49^, and directly using asymmetric vesicles to form SLBs^50^. While these methods enable the generation of asym-SLBs, they are often limited to specific lipid types for asymmetric incorporation, and they do not offer precise control over the asymmetric distribution of lipids between the two leaflets. Recently, an elegant technique has been developed to generate asym-SLBs for PIP_2_, in which additional vector small unilamellar vesicles (SUVs) containing PIP_2_ were introduced to dope the top leaflet of pre-formed SLBs with PIP_2_, although the underlying formation mechanism remains to be identified^35^. To date, there is a lack of robust methods to generate SLBs with asymmetric distribution of PIP lipids on the top leaflet of the SLBs, facilitating quantitative studies of lipid-protein interactions.

In this study, we propose a new method for preparing asym-SLBs enriched in PIP_2_ exclusively in the top leaflet of the SLBs. Inspired by a recently developed method involving PIP_2_-containing vector SUVs^35^, our approach takes advantage of lipids with high critical micelle concentrations (CMC). At concentrations below the CMC, lipids do not form micelles but remain as monomers. Specifically, in our method, PIP_2_ is introduced in solution at concentrations below its CMC, enabling its spontaneous insertion into pre-formed symmetric SLBs. This strategy is analogous to previous reports of spontaneous insertions of short chain ceramides and single-chain lysolipids into the outer leaflet of liposomes^51,52^.

Our method is simple and robustly reproducible. It allows control over the concentration of the lipids of interest incorporated into asym-SLBs. The asym-SLBs allow us to study the dynamics of the lipids of interest free from any interaction with the substrate. We quantitatively characterized membrane asymmetry and fluidity using fluorescently labelled PIP_2_, TopFluor-PIP_2_ (TF-PIP_2_). Our method allows the amount of TF-PIP_2_ incorporation to be fine-tuned. We further demonstrated the utility of asymmetrically incorporating PIP_2_ into SLBs by recruiting the PIP_2_-associated actin-membrane linker ezrin, and myosin I motor, myosin 1b, which glides single actin filaments on the PIP_2_-asym-SLBs. Taken together, the asym-SLBs developed in this study provide a robust experimental model system for future quantitative studies of lipid dynamics and lipid-protein interactions in vitro.

## Materials and Methods

### Reagents

1,2-Dioleoyl-sn-glycero-3-Phosphatidylcholine (DOPC, 850375), 1,2-dioleoyl-sn-glycero-3-phospho-L-serine (DOPS, 840035), L-α-phosphatidylinositol-4,5-bisphosphate (PIP_2_, Brain, Porcine, 840046P), and 1-oleoyl-2-[6-[4-(dipyrrometheneboron difluoride)butanoyl]amino]-hexanoyl-sn-glycero-3-phosphoinositol-4,5-bisphosphate (TF-PIP_2_, 810184P) were purchased from Avanti Polar Lipids. Alexa Fluor 488 C5-Maleimide (AX488) and Alexa Fluor 647 (AX647) phalloidin were purchased from Invitrogen. Creatinine phosphate was purchased from Roche (10621714001) and creatinine phosphate kinase from Sigma-Aldrich (C3755-3.5KU). Other reagents were purchased from Sigma-Aldrich.

### Buffer

The buffer used to prepare PIP_2_ solutions and to wash the SLBs, referred to as the working buffer, contains 0.8 mM Sodium Phosphate, 15.3 mM Sodium Chloride, 0.3 mM Potassium Chloride, 0.2 mM Potassium Phosphate and 1 mM EDTA, pH 7.5. For PIP_2_ clustering experiments, the 1 mM EDTA is replaced by 1 mM CaCl_2_. MilliQ (MQ) water was used in all buffer preparations.

### Preparation of SUVs

The desired amounts of lipids were mixed in CHCl_3_ and then dried with N_2_, followed by drying in a vacuum chamber overnight to ensure that the CHCl_3_ was completely evaporated. The dried lipid film was hydrated with MQ H_2_O for lipid mixtures without PIP_2_, or the working buffer for lipid mixtures containing PIP_2_, at a lipid concentration of 1.5 mg/mL and incubated at room temperature for 30 min to form vesicles. The resulting vesicle suspension was vortexed, followed by sonication in a bath sonicator (Fisher Scientific, ref. 10611983, at maximum power), twice for 30 min each, at room temperature, to generate SUVs. The resulting SUVs were then centrifuged at 20000g for 1 hour at 4°C to remove any lipid debris. The SUV supernatant was collected and stored at 4°C for a maximum of two weeks.

### Preparation of PIP_2_ solutions

The desired amount of PIP_2_ in CHCl_3_ was dried with N_2_, followed by drying in a vacuum chamber for a few hours or overnight to ensure that the CHCl_3_ was completely evaporated. The resulting lipid film was resuspended in the working buffer to obtain a final lipid concentration of 10 µM for PIP_2_ and TF-PIP_2_. The lipid solution was either stored at 4°C for a maximum of two weeks or at -20°C for several months.

### Preparation of PIP_2_-asym-SLBs

Glass coverslips were first cleaned by sonication in the following solutions for 20 min each: H_2_O, Acetone, H_2_O, 1M KOH and 5 times with H_2_O. The coverslips were dried with N_2_ and stored under vacuum before use. The experimental chamber was assembled from two cleaned coverslips glued together using double sided tapes, with the bottom coverslip plasma cleaned for 3 min to make it hydrophilic. SUVs composed of DOPC:DOPS at 9:1 molar ratio were prepared following the above-mentioned method. The chamber was filled with SUVs at 0.5 mg/mL, supplemented with 1 mM CaCl_2_, followed by incubation at room temperature for 30 min to form SLBs. The resulting SLB was rinsed with 200 µL working buffer (∼10 times the chamber volume). Then, the chamber was filled with PIP_2_ or TF-PIP_2_ solution for 10 min, followed by rinsing with 100 µL working buffer. The asym-SLB was used immediately for experiments.

### Preparation of sym-SLBs containing TF-PIP_2_

SUVs composed of DOPC:DOPS:TF-PIP_2_ at 85:10:5 molar ratio were prepared following the above procedure, using the working buffer to rehydrate lipid films. An experimental chamber was filled with SUVs at 0.5 mg/mL, supplemented with 3 mM CaCl_2_, followed by incubation at room temperature for 30 min to form SLBs. The resulting SLB was rinsed with 200 µL working buffer (∼10 times the chamber volume). The SLB was used immediately for experiments.

### FRAP experiments and data analysis

Confocal imaging was performed on an inverted microscope (Eclipse Ti-E, Nikon) equipped with a spinning disk CSU-X1 (Yokogawa), a sCmos camera Prime 95B (Photometrics) and a 60X objective (NA 1.40, oil, Nikon). A laser bench of four lines (405nm, 491nm, 561nm, 642nm; Gataca Systems) was used for FRAP and acquisition at 491 nm. The fluorescent intensities at each time points were obtained using Metamorph software (Gataca Systems). For each image, the intensity of the bleached area was corrected with the intensity of an unbleached area, far from the bleached region, to account for photobleaching and potential variation of the intensity overtime. For each FRAP experiment, we performed FRAP with four bleaching areas with diameters of 9.2 µm, 16.5 µm, 23.8 µm and 31.2 µm. We performed three FRAP measurements for each area sizes. The analysis of the FRAP curves and the extraction of characteristic time and immobile fraction were performed as previously described^53^. The diffusion coefficient was obtained by fitting the diffusion coefficient using the following equation: 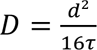, where the values of *d* and *τ* obtained from each individual FRAP measurement, and *D* is the diffusion coefficient, *d* is the diameter of the bleached area, and *τ* is the characteristic time of the FRAP recovery curve. (See supplementary figure 1)

### Quantification of TF-PIP_2_ incorporation into asym-SLBs

To quantify the incorporation of TF-PIP_2_ into the top leaflet of asym-SLBs, we compared their fluorescent intensity with that of sym-SLBs. Sym-SLBs served as a reference because their TF-PIP_2_ content is directly determined by the lipid composition of the SUVs used for their preparation. Specifically, sym-SLBs were formed from SUVs containing DOPC supplemented with 10 mol% DOPS, and with 0.1, 1, 2, 3 or 4 mol% TF-PIP_2_. For each confocal image of SLBs, the mean fluorescent intensity was calculated within a 200 x 200 pixel region. These data were used to generate a calibration curve, yielding the slope *m_sym-SLB,_* (intensity units per mol%) (Fig. S2). All imaging and analysis were performed under identical microscope settings, including laser power and detector gain, for both sym-SLBs and asym-SLBs. The mole fration of TF-PIP_2_ in the top leaflet of aysm-SLBs was then calculated as 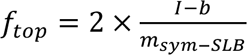, where *I* is the measured intensity of asym-SLBs and *b* is the background intensity. The factor of 2 accounts for the fact that in sym-SLBs TF-PIP_2_ is present in both leaflets, whereas in asym-SLBs it is present only in the top leaflet.

### Determination of the critical micelle concentration (CMC) of TF-PIP_2_

Solutions of TF-PIP_2_ in the working buffer are prepared at TF-PIP_2_ concentrations ranging from 1 µM to 2 mM. The principle of the CMC determination is that monomers of TF-PIP_2_ are not quenched and are thus visible in fluorescence spectrometry, whereas TF-PIP_2_ in micelles is self-quenched. When TF-PIP_2_ concentrations are increased below the CMC, the fluorescence intensities are thus expected to increase proportionally to the concentration but the fluorescence intensity will plateau as soon as the CMC is reached and additional TF-PIP_2_ is incorporated in micelles. The fluorescence intensities of TF-PIP_2_ were measured with a Tecan microplate spectrophotometer.

### Thermodynamic model of TF-PIP_2_ incorporation in asym-SLBs

The principle of the thermodynamic model is the thermal equilibrium between the three different forms of TF-PIP_2_ (monomer in solution, in a micelle in solution, inserted in the SLB):

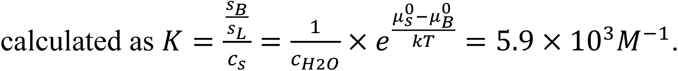

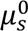, 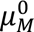, and 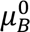 are the standard chemical potentials of TF-PIP_2_ in solution (*s*), micelles (*M*), and asym-SLBs (*B*), respectively. 𝑐_𝑠_ and 𝑐_𝑀_ are the bulk concentrations of TF-PIP_2_ in solution and in micelles, *c_H_*_20_ = 55.6 𝑀 is the water concentration. 𝑠_𝐵_ and 𝑠_𝐿_ are the surface densities of TF-PIP_2_ and lipids in asym-SLBs, respectively (*S_L_* = 1.3 × 10^18^ molecules/m^2^ if the surface per lipid is 0.7 nm^2^). N is the number of TF-PIP_2_ molecules per micelle; this number is not known but its exact value has no impact here provided that the monomer concentration remains below CMC and that N>∼10.

The total TF-PIP_2_ concentration is computed by taking into account the TF-PIP_2_ fraction in the SLB, knowing that our sample has a volume of 20 µL and our SLB covers 2.6 cm^2^.

The model must reproduce: (1) the proportionality between the proportion of TF-PIP_2_ in asym-SLB (%) and the total TF-PIP_2_ concentration (µM) when the proportion is below 5% (no self-quenching), and (2) the TF-PIP_2_ CMC = 290 µM. A good fit is obtained with: 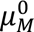 = 0 𝑘𝑇 (our reference), 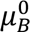 = −0.6 𝑘𝑇 and 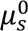 = 12.2 𝑘𝑇 (Fig. 3, blue line). From these standard chemical potentials, the TF-PIP_2_ partition coefficient between the membrane and the solution can be calculated as 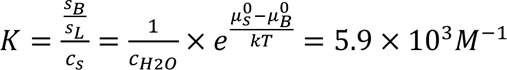.

**Table 1.**
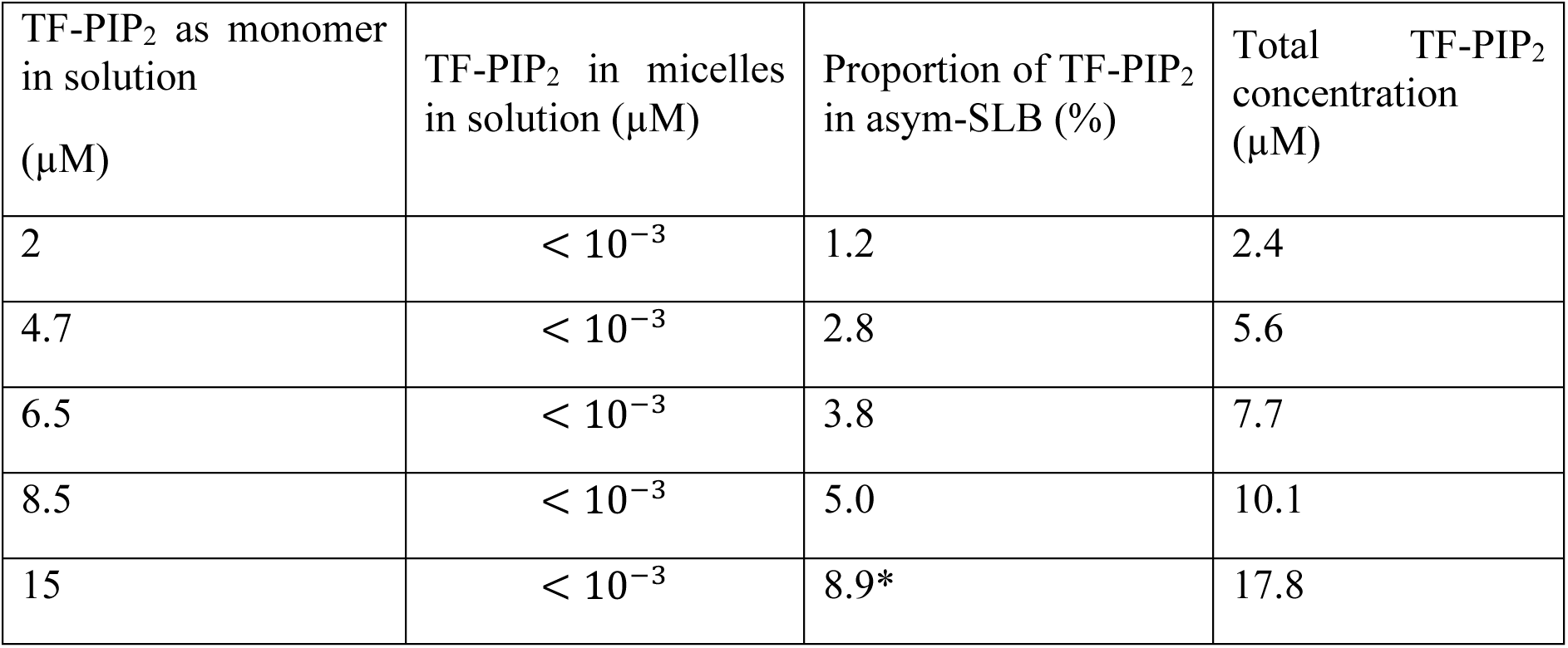
Thermodynamic model prediction of TF-PIP_2_ in asym-SLBs. *at the bulk concentration of 15 μM, TF-PIP_2_ signal is quenched.

| TF-PIP<sub>2</sub> as monomer in solution (µM) | TF-PIP<sub>2</sub> in micelles in solution (µM) | Proportion of TF-PIP<sub>2</sub> in asym-SLB (%) | Total TF-PIP<sub>2</sub> concentration (µM) |
| --- | --- | --- | --- |
| 2 | < 10<sup>-3</sup> | 1.2 | 2.4 |
| 4.7 | < 10<sup>-3</sup> | 2.8 | 5.6 |
| 6.5 | < 10<sup>-3</sup> | 3.8 | 7.7 |
| 8.5 | < 10<sup>-3</sup> | 5.0 | 10.1 |
| 15 | < 10<sup>-3</sup> | 8.9* | 17.8 |

### Measurement of TF-PIP_2_ self-quenching

To monitor the conditions of TF-PIP_2_ self-quenching, we prepared liposomes in water at 1 mM POPC final concentration with various TF-PIP_2_ concentrations, so that the proportion of TF-PIP_2_ in liposome membranes varies from 0% to 10%. To obtain the right TF-PIP_2_ proportion, we have taken into account the partition of TF-PIP_2_ between membranes and solution according to the partition coefficient we have determined (𝐾 = 5.9 × 10^3^𝑀^−1^), which leads to adding ∼20% more TF-PIP_2_ than if it is fully embedded in liposome membranes. The fluorescence intensities of the liposome samples were measured with a Tecan microplate spectrophotometer.

As shown in Fig. S3, we observed that the fluorescence intensities of TF-PIP_2_ increase proportionally to the TF-PIP_2_ percentage in the liposome membrane up to 5% TF-PIP_2_. Above 5%, the fluorescence intensity remains approximately constant, which can be explained by TF-PIP_2_ self-quenching.

### Myosin 1b, actin and ezrin purification

Human myosin 1b was purified from HEK293-Flp-In cells following the method described previously^54^. Myo1b was stored in the following buffer: 30 mM Hepes pH 7.5, 150 mM KCl, 4 mM MgCl_2_, 1 mM EGTA, 0.1% Methylcellulose, 4 mM Na_2_ATP, and 1 mM DTT. Muscle actin was purified from rabbit muscle and isolated in monomeric form in actin buffer (5 mM Tris-HCl, pH 7.8, 0.1 mM CaCl_2_, 0.2 mM ATP, 1 mM DTT and 0.01% NaN_3_)^55^. Phosphomimetic ezrin T567D (ezrin TD) was purified and labelled with AX488 as previously described^56^. EzrinTD was stored in 20 mM Tris pH 7.4, 50 mM KCl, 0.1 mM EDTA, 2 mM β-Mercaptoethanol, and supplemented with 0.1% methylcellulose.

### Preparation and analysis of ezrin-actin binding assays

To evaluate ezrin binding and actin recruitment on PIP₂-asym-SLBs, base SLBs were prepared by the fusion of SUVs composed of DOPC: DOPS at a 9:1 molar ratio and supplemented with 0.5 mol% Texas Red-DHPE for membrane visualization. SUVs were prepared as described above and introduced into the chamber at a final lipid concentration of 0.5 mg/mL. The chamber was incubated with the SUVs for 30 min at room temperature, protected from light. SLBs were then gently washed with 200 µL of the working buffer to remove excess vesicles.

After the formation of the base SLB, 20 µL of a 10 µM PIP₂ solution was added directly to the chamber, followed by incubating with the SLB for at least 10 min. The chamber was subsequently washed with ∼100 µL of the working buffer. For ezrin recruitment assays, phosphomimetic ezrin T567D (ezrin TD) labeled with AX488 was prepared at 10 µM in 20 mM HEPES pH 7.4, 50 mM KCl, and 0.1 mM EDTA. The protein solution (20 µL) was added to the chamber and incubated for 10 min. Next, pre-polymerized actin filaments labeled with AX647-tagged phalloidin were introduced in the chamber, followed by incubating for 10 min at room temperature. The chamber was sealed and prepared for imaging. Control experiments were performed using base SLBs, without PIP_2_ incorporation.

Actin filaments were prepared by polymerizing globular actin monomers at 5 µM, then diluted to 1 µM in the actin polymerization buffer (5 mM Tris HCl pH 8.0, 3 mM MgCl_2_, 0.2 mM EGTA, 1 mM DABCO, 10 mM DTT, 100 mM KCl, 2 mM ATP), supplemented with AX647-taggd phalloidin at 330 nM.

Samples were imaged using a spinning-disk confocal microscope: Inverted Eclipse Ti-E (Nikon) equipped with a CSU-X1 spinning disk (Yokogawa), controlled by Metamorph software (Gataca Systems), and a 100x oil objective. Imaging was performed using appropriate laser lines and filters for Texas Red, AX488, and AX647. The pixel-averaged intensity of ezrin fluorescence was calculated from a 500 × 500 pixel-sizes for each image.

### Preparation and analysis of motor motility assays

The actin motility assays were performed and analyzed following the method described previously^57^. Briefly, myo1b at 600 nM was incubated with PIP_2_-asym-SLBs for 30 min at room temperature. Then, a 50 nM solution of pre-polymerized actin filaments (in the actin polymerization buffer, supplemented with 0.3% methylcellulose and the ATP regeneration system) stabilized with AX647 phalloidin at 165 nM was introduced into the experimental chambers. The ATP regeneration system is composed of 2 mM Na_2_ATP, 2 mM MgCl_2_, 20 mM Creatinine phosphate and 3.5 U/mL of Creatinine phosphate kinase.

The samples were observed using TIRF microscopy (Eclipse Ti inverted microscope, 100X TIRF objective, Quantem512SC camera). Kymographs of gliding actin filaments were generated using the ImageJ plug-in KymoToolBox.

## Results and Discussions

### The generation of asymmetric SLBs with (PIP2-asym-SLBs)

A wide range of CMC values of PIP_2_ has been reported. For unlabeled PIP_2_, the CMC is on the order of a few μM in 100 mM NaCl, 30 μM in 50 mM PIPES or Tris pH 7, and 200 μM in water^58–60^. For BODIPY® TMR PIP_2_ in water, the CMC value was estimated to be up to 500 μM^58–60^. We hypothesized that dissolving PIP_2_ at a relatively high concentration in solution, but below its CMC, could facilitate its insertion into the top leaflet of pre-formed SLBs. This spontaneous lipid insertion is due to the less favorable hydrophobic interaction of the fatty acid chains of PIP_2_ with the solution than with the lipid membrane. We first determined the CMC of TF-PIP_2_ in the Working buffer (0.8 mM Na₃PO₄, 15.3 mM NaCl, 0.3 mM KCl, 0.2 mM KH₂PO₄, and 1 mM EDTA, pH 7.5), as used previously^35^. We prepared TF-PIP_2_ solutions at concentrations ranging from 1 µM to 2 mM. The principle of CMC determination is that monomeric TF-PIP_2_ is not quenched and its fluorescent signal is detectable in fluorescence spectrometry, while TF-PIP_2_ in micelles is self-quenched. Thus, TF-PIP_2_ fluorescence intensity increases linearly with concentrations below the CMC, then plateaus once the CMC is reached due to micelle formation. As shown in Fig. 1A, the CMC of TF-PIP_2_ was determined to be 290 μM in the working buffer.

**Figure 1.**
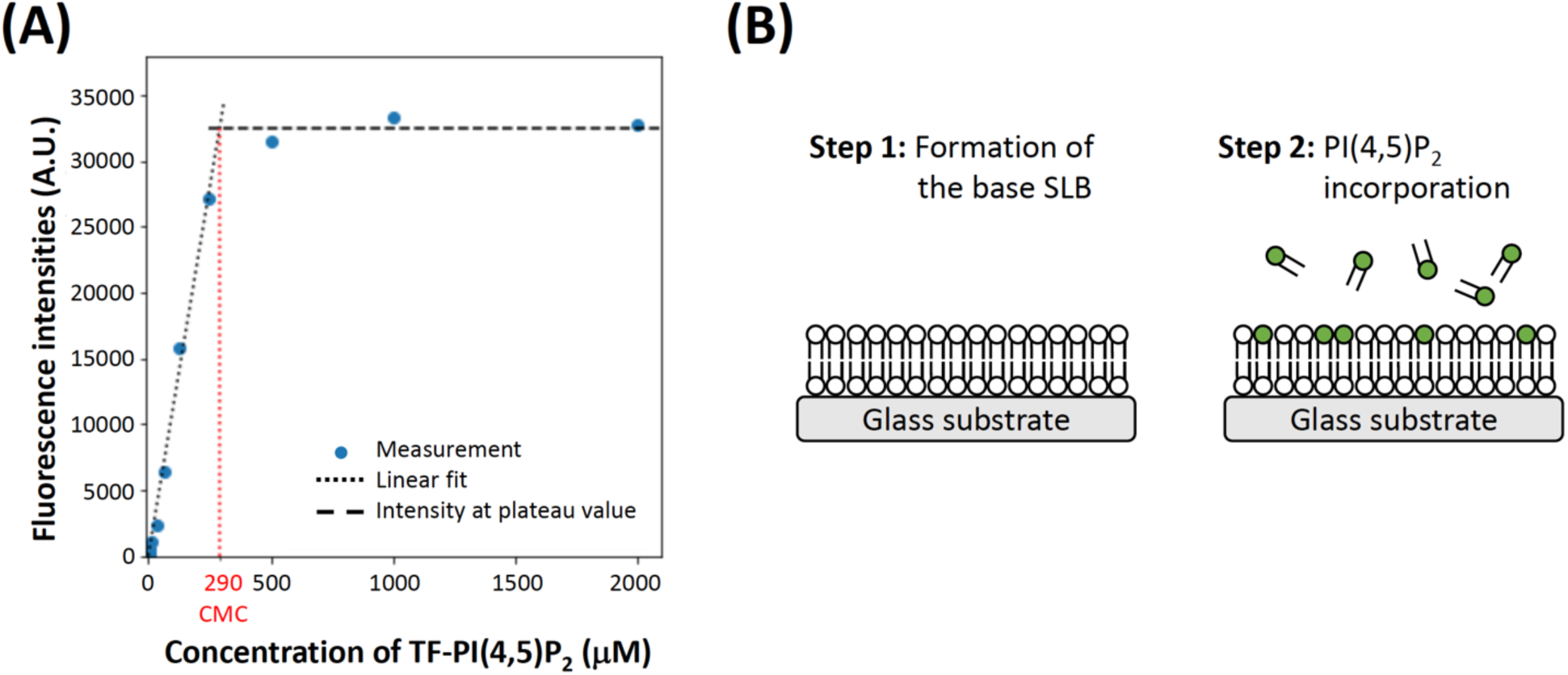
Proposed mechanism for PIP_2_ asym-SLB formation. (A) Determination of the CMC of TF-PIP_2_ based on fluorescence intensities measurements (blue symbols. See Materials and Methods for details.). Dotted line is the linear fit of the measurements. Dashed line represents the plateau value of the intensities at high TF-PIP_2_ concentration. The CMC, 290 μM, is indicated in red. (B) Schematic representation of PIP_2_ asym-SLB formation. Step 1: Formation of a base SLBs composed of DOPC and DOPS via SUV fusion on a glass substrate. Step 2: Introduction of a PIP_2_ solution at a concentration below CMC, enabling its spontaneous insertion into the top leaflet of the SLB and resulting in an asymmetric TF-PIP_2_ distribution. Green-headed lipids represent PIP_2_.

To test our hypothesis of TF-PIP_2_ spontaneous insertion into SLBs, we first prepared a SLB containing DOPC and DOPS (9:1 molar ratio) on a glass substrate using the well-established SUV fusion method^61^. Then, a solution of TF-PIP_2_ at a concentration below its CMC, at 5 μM, was introduced into the experimental chamber. We incubated the base SLB with TF-PIP_2_ for 10 min, allowing it to insert into the top leaflet of the base SLB (Fig. 1B). The resulting asym-SLB was rinsed with the working buffer to remove excess TF-PIP_2_ remaining in the solution. Using 5 µM of TF-PIP_2_ in solution produced asym-SLBs with relatively homogenous fluorescence, demonstrating the proper incorporation of TF-PIP_2_ on the SLBs (Fig. 2A *Top panel*).

**Figure 2.**
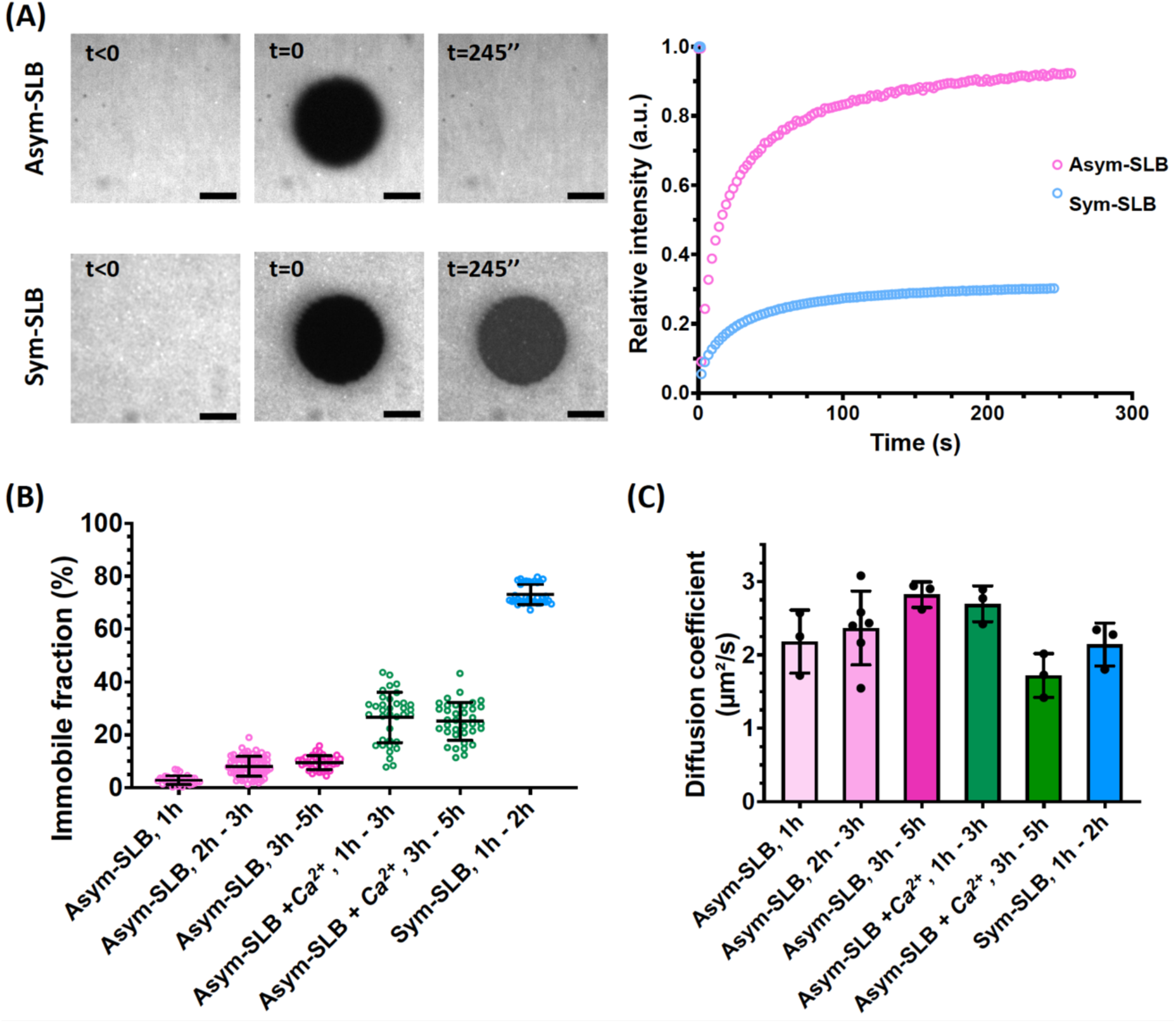
Quantitative comparisons of sym- and asym-SLBs for TF-PIP_2_, and the effect of Ca^2+^ in asym-SLBs. **(A)** Left: Representative confocal images of FRAP experiments on TF-PIP_2_ in sym- and asym-SLBs before bleaching (t<0), immediately after bleaching (t=0) and after recovery (t=245 sec). Scale bars, 10µm. Right: Representative relative fluorescent intensities of the bleached area overtime. By analyzing the FRAP curves, we found the recovery of ∼98% and ∼30% and the diffusion coefficient of 2 μm^2^/sec and 1.5 μm^2^/sec of TF-PIP_2_ in the asym-SLB (Magenta circles) and sym-SLB (Cyan circles), respectively. **(B)** Quantification of the immobile fractions of TF-PIP_2_ in FRAP experiments performed within durations of one hour (1h), 2h-3h and 3h-5h after bilayer formation. N = 3 independent experiments, except “asym-SLB 2h-3h” with N = 6, for each condition, in which n = 34, 71, 36, 36, 36, 36 FRAP measurements (from left to right). The immobile fractions were calculated by data fitting to the FRAP curves. **(C)** Diffusion coefficients of TF-PIP_2_ in asym- and sym-SLB. For the asym-SLBs, TF-PIP_2_ bulk concentration was 5 μM. Each data point represents one independent FRAP experiment. Within each experiment, there were 4 different bleaching area sizes and 3 FRAP measurements were performed for each size. Thus, a total of 12 FRAP measurements in one FRAP experiment. Diffusion coefficients for each experiment were obtained from the linear fit of bleached areas to their recovery characteristic times (as shown in Fig. S1). The error bar represents the standard deviation. Statistic test, Mann-Whitney test, for conditions of Asym-SLB 3h-5h and Asym-SLB + Ca^2+^ 3h-5h, p = 0.1. In (B) and (C), for experiments with Ca^2+^, Asym-SLB + Ca^2+^, in the working buffer, 1 mM EDTA was replaced by 1 mM CaCl_2_.

**Figure 3.**
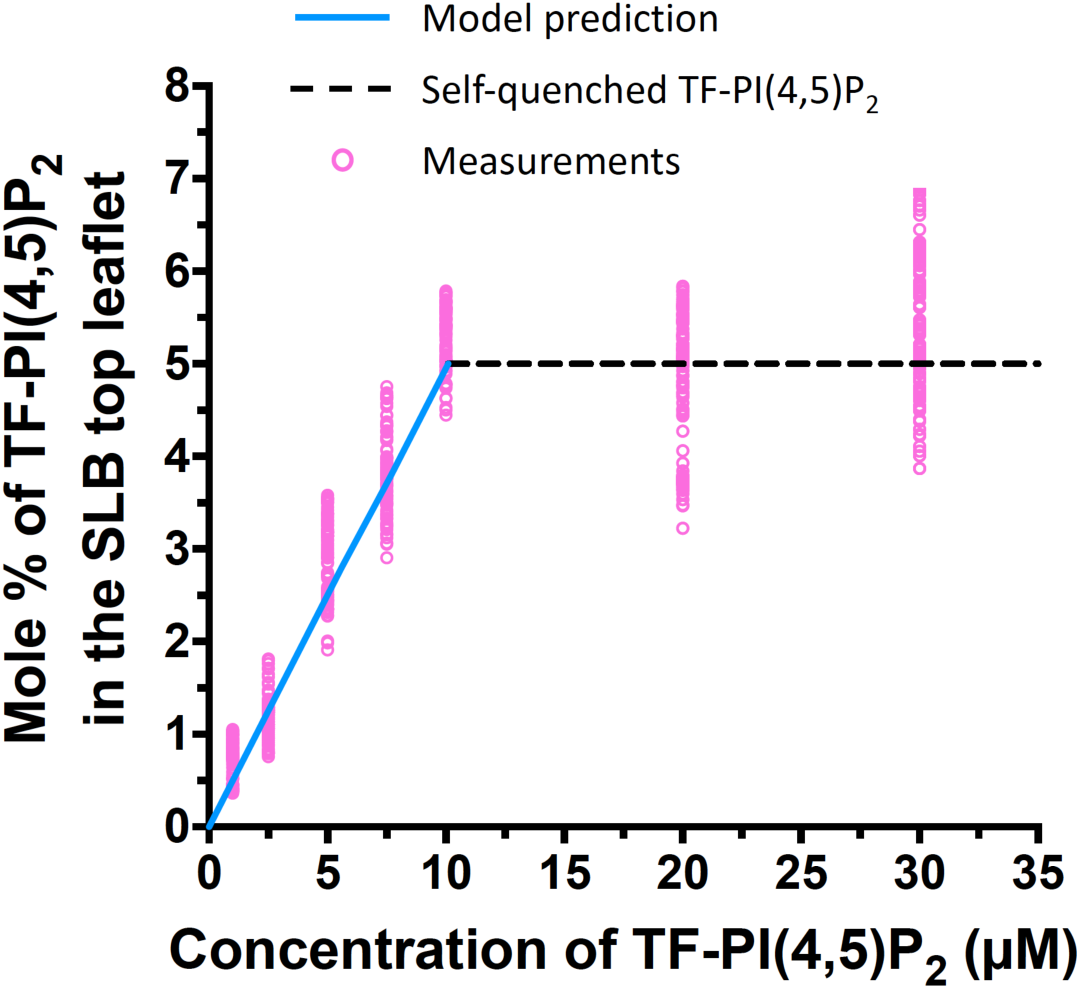
Quantification of TF-PIP_2_ incorporation into asym-SLBs. Quantification of the molar percentages of TF-PIP_2_ in the top leaflet of asym-SLB generated by using different bulk concentrations of TF-PIP_2_. The blue line is the prediction of the thermodynamic model for all the data points < 10µM, slope = 0.53. n = 568 measurements, N = 7 independent experiments.

### Quantitative characterization of PIP2-asym-SLBs

To quantitatively characterize the resulting TF-PIP_2_-asym-SLBs, we performed Fluorescence Recovery After Photobleaching (FRAP) experiments to measure the diffusion coefficients of TF-PIP_2_ and its immobile fractions within the asym-SLBs (Fig. 2A, Asym-SLB). For freshly prepared asym-SLBs, within 1 hour after preparation, we found the immobile fractions of 2.8 ± 1.7 % (Fig. 2B, Asym-SLB, 1h). By analyzing the FRAP curves (Fig. S1), we found that the diffusion coefficients of TF-PIP_2_ in these asym-SLBs were around 2 ± 0.4 μm^2^/sec (Fig. 2C, Asym-SLB, 1h), which are comparable to previously reported values for SLBs^34–36^. The initial residual immobile fraction in TF-PIP_2_ asym-SLBs may be due to the presence of unfused SUVs in the pre-formed SLBs, as previously shown by scanning electron microscopy^62^. Notably, by unzipping lipid monolayers from SLBs, a recent study demonstrated that PIP_2_ can translocate from the top to the bottom leaflet at the edges of bilayer defects in sym-SLBs, contributing to the immobilized fraction^36^. Moreover, it was reported that the translocation of PIP_2_ from the top to the bottom leaflet at bilayer defects sites results in a progressive increase in the PIP_2_ immobile fraction over time^36^. Here, we compared the immobile fraction of a freshly prepared asym-SLB with those of asym-SLBs a few hours after formation. Indeed, we found the mean immobile fractions increased after formation, up to 9% after 5 hours (Fig. 2B, Asym-SLB, 2h-3h and 3h-5h), while there is no significant difference for the diffusion coefficients, ranging between 1.5 – 3 μm^2^/sec (Fig. 2C, Asym-SLB, pink bars, *p*=0.1 one way ANOVA test). These results are consistent with the previous report^36^, and suggest that, under our experimental conditions, TF-PIP_2_ in asym-SLBs undergoes a modest degree of translocation over the timescale of the experiments.

We compared the immobile fractions and diffusion coefficients of TF-PIP_2_-asym-SLBs with those measured in sym-SLBs containing TF-PIP_2_. We prepared the symmetric SLBs using SUVs composed of DOPC:DOPS:TF-PIP_2_ at 88:10:2 molar ratio. Given the known tendency of PIP_2_ to remain immobile when in contact with a glass substrate^31,35,36^, the measured immobile fraction of PIP_2_ serves as an indicator of its asymmetric distribution in the two leaflets of the SLB. The time series of the FRAP images showed a clear difference between the sym- and asym-SLBs (Fig. 2A).

While the asym-SLB exhibited an immobile fraction of less than 20% (Fig. 2B, Asym-SLB, pink data points), the sym-SLB showed much less fluorescence recovery, reflecting a higher immobile fraction of 75% on average (Fig. 2B, Sym-SLB, blue data points). In the sym-SLB, the higher immobile fraction of TF-PIP_2_ can be partly attributed to the fact that 50% of the TF-PIP is in direct contact with the glass substrate, resulting in its immobilization, as previously observed^33–36,63^. For the mobile fraction of TF-PIP_2_, we found similar diffusion coefficients in both asym- and sym-SLBs (Fig. 2C, compare pink and blue bars). This similarity probably arises from the fact that mobile PIP_2_ lipids are only present in the top leaflet of both asym- and sym-SLBs. Overall, we showed here that the proposed method based on the spontaneous insertion of PIP_2_ into pre-formed sym-SLBs allows us to generate asym-SLBs with preserved PIP_2_ fluidity.

Divalent ions such as calcium and magnesium are known to cluster PIP_2_ via electrostatic interactions, which can affect the immobile fraction and potentially alter the diffusion of PIP_2_^12–14,64,65^. To evaluate how well PIP_2_ in the asym-SLBs retains its expected properties, such as its ability to diffuse or form clusters in the presence Ca^2+^ ions, we quantified the effect of Ca^2+^ on the diffusion and immobile fractions of TF-PIP_2_. By performing FRAP experiments, we observed an increase in the immobile fraction of TF-PIP_2_ in the presence of 1 mM of Ca^2+^ ions compared to that without Ca^2+^ ions (Fig. 2B, green data points). The increased immobile fraction indicates PIP_2_ molecules were clustered by Ca^2+^ ions. On the other hand, the diffusion coefficient of mobile TF-PIP_2_ was only slightly affected by the presence of Ca^2+^ ions (Fig. 2C, green bars). This observation is consistent with a previous study using FCS reported a slight decrease in the diffusion of TF-PIP_2_ monomers and TF-PIP_2_ clustering in GUVs due to the presence of Ca^2+^ ions^11^. Our results indicated that in the presence of Ca^2+^, the freely diffusing TF-PIP_2_ was not significantly affected by the presence of TF-PIP_2_ clusters and possibly the kinetics of clustering and de-clustering is much slower than the timescale used for measuring the FRAP recoveries.

### Control of the PIP2 amount incorporated into the top leaflet of pre-formed SLBs

To quantify the amount of PIP_2_ incorporated into the top leaflet of the SLBs, we measured the fluorescent intensity of TF-PIP_2_ in asym-SLBs. We calibrated our measurements using TF-PIP_2_ fluorescence measurements on sym-SLBs since the concentrations of TF-PIP_2_ incorporated into the SUVs used to generate the sym-SLBs is known (see Method section for details). We found that the amount of TF-PIP_2_ incorporated into the top leaflet of asym-SLBs was directly proportional to the bulk concentration of TF-PIP_2_, up to 10 μM of TF-PIP_2_ (Fig. 3). The use of TF-PIP_2_ at bulk concentrations ranging from 0 to 10 µM resulted in 0 to 5 mol% TF-PIP_2_ in asym-SLBs (Fig. 3).

We observed that beyond 10 µM of TF-PIP_2_, the intensity of TF-PIP_2_ did not increase significantly. This observation suggests that either the asym-SLBs were saturated with TF-PIP_2_, or the concentration of TF-PIP_2_ was high enough for the TF dyes to quench each other^12,66^. To test for self-quenching, we prepared vesicles containing 0 mol% to 10 mol% TF-PIP_2_. We observed self-quenching when the TF-PIP_2_ fraction above 5 mol% (Fig. S3), supporting the hypothesis that fluorescence saturation at higher concentrations is due, at least in part, to quenching effects.

To quantitatively explain our observations of TF-PIP_2_ incorporation into SLBs or micelles, we implemented a thermodynamic model of the equilibrium between monomeric and micellar TF-PIP_2_ in solution and TF-PIP_2_ inserted into the SLB (see method section). Using our experimentally observed proportionality between TF-PIP_2_ incorporation in SLBs and bulk concentration below 5 mol% of TF-PIP_2_, we used our thermodynamic model to calculate the TF-PIP_2_ partition coefficient between the membrane and the solution to be 5.9 × 10^3^𝑀^−1^.

### The activity of PIP2-associated ezrin and molecular motor myosin 1b on PIP2-asym-SLBs

Actin-membrane linkers often contain domains that specifically interact with PIP_2_. To further validate the use of PIP_2_-asym-SLBs, we examined the recruitment of ezrin, a well-characterized actin-membrane linker^18^. Ezrin contains an N-terminal FREM (band 4.1, ezrin, radixin, moesin) domain that binds specifically to PIP_2_ and a C-terminal ERM associated domain that interacts with actin filaments. We assessed both ezrin recruitment to PIP_2_-asym-SLBs and its ability of anchoring actin filaments to the bilayers. We incubated PIP_2_-asym-SLBs with 10 μM AX488-labelled phosphomimetic ezrin T567D (ezrin TD), which adopts an open conformation that allows simultaneous interactions with PIP_2_ and actin. This was followed by the addition of 1 μM AX647-phalloidin-labelled actin filaments. In the absence of PIP_2_, neither ezrinTD nor actin bound to the SLBs (Fig. 4A). However, when PIP_2_ was present, both ezrin and actin filaments were clearly recruited to the membrane (Fig. 4, A and B). Additionally, we investigated the activity of the class I myosin motor, myo1b, which contains a C-terminal tail pleckstrin homology (PH) domain that binds PIP_2_ and an N-termial motor domain that binds actin filaments and generates force^67^. We incubated PIP_2_-asym-SLBs with 1 µM myo1b and 50 nM of AX647-phalloidin labelled actin filaments to perform motility assays^68^. We observed the gliding of the actin filaments on asym-SLB containing TF-PIP_2_ or non-fluorescently labelled PIP_2_, showing myo1b activity can be reconstituted in this system (Fig. 4C). The resulting actin gliding velocities are 57 nm/sec for TF-PIP_2_ and 62 nm/sec for non-fluorescently labelled PIP_2_, both comparable to those previously measured for myo1b on PIP_2_-sym-SLBs (Fig. 4D)^57^.

**Figure 4.**
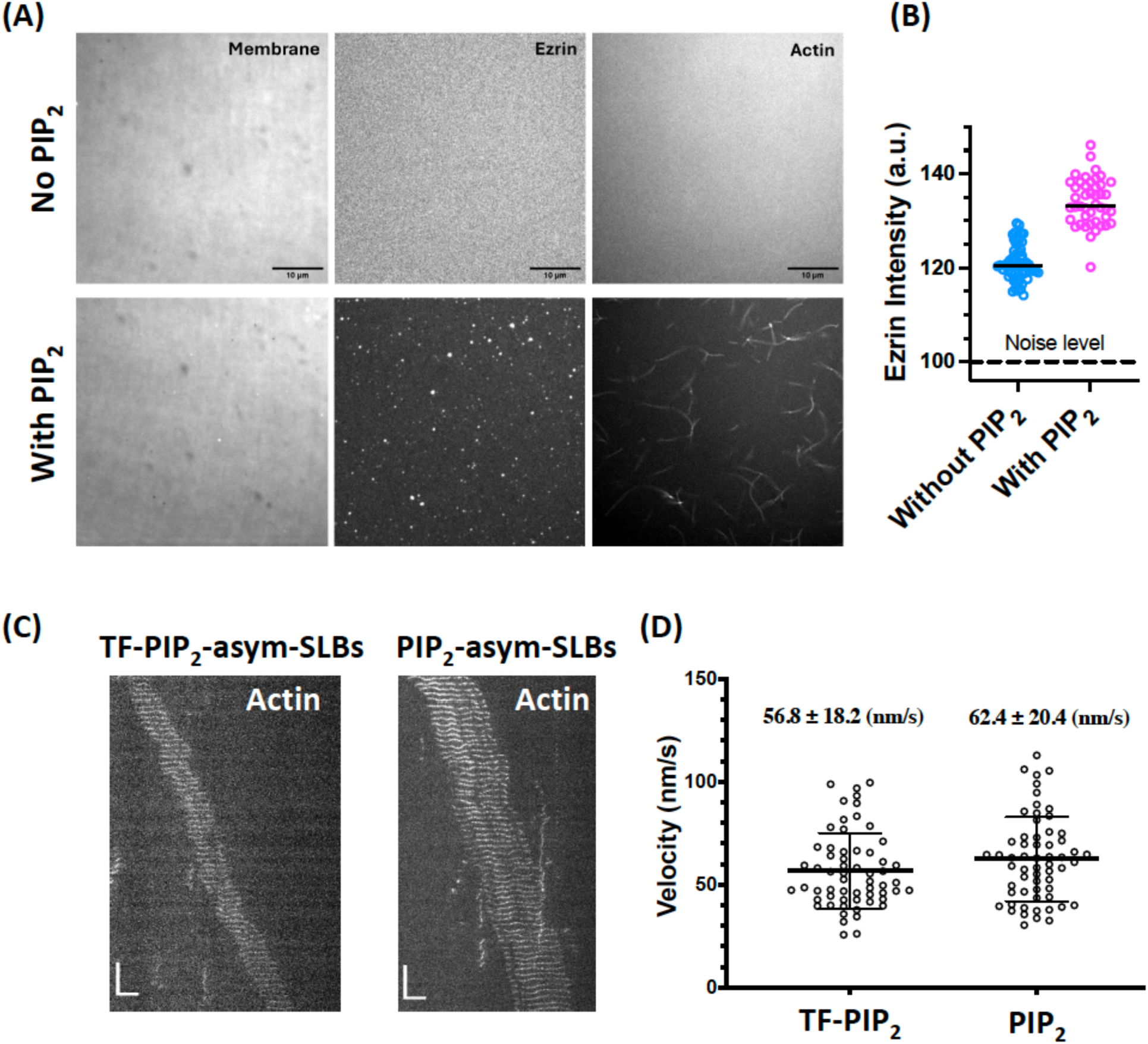
**Actin filament recruitments on PIP_2_-asym-SLBs by actin-membrane linkers, ezrin and myo1 motors**. **(A)** Representative confocal images of AX488 labelled ezrinTD and AX647 phalloidin-labelled actin filaments on SLBs without PIP_2_ (Top) and PIP_2_-asym-SLBs (Bottom). From Left to Right panel: Membrane, Ezrin and Actin. Base SLBs were composed of DOPC: DOPS at a 9:1 molar ratio and supplemented with 0.5 mol% Texas Red-DHPE for membrane visualization. The asym-SLBs were generated by incubating the base SLBs with 10 μM PIP_2_ solution. Scale bars, 10 µm. **(B)** Quantification of ezrin intensity on SLBs without PIP_2_ and PIP_2_-asym-SLBs. N = 3 independent experiments. Analyzed images: 60 for SLBs without PIP_2_ and 44 for PIP_2_-asym-SLBs. **(C)** Representative kymographs of actin filament gliding by myo1b bound on asym-SLBs with PIP_2_ or TF-PIP_2_. Base SLBs were incubated with 5 μM PIP_2_ or TF-PIP_2_ solutions. Horizontal scale bars, 5 µm, Vertical scale bars, 1 min. Fluorescent singles were AX647 phalloidin labelled actin filaments. **(D)** Velocities of single actin filament gliding by myo1b bound on asym-SLBs with TF-PIP_2_ or PIP_2_. The mean velocities and the corresponding standard deviations of myo1b-driven actin filament gliding are indicated in the figure. Welch’s t test, p = 0.1119. For TF-PIP_2_, n=62 filaments measured and N=3 independent experiments. For PIP_2_, n=60 filaments measured, N=2 independent experiments.

## Conclusion

In this study, we introduced a simple two-step method to generate supported bilayers containing leaflets with different lipid compositions. We demonstrated the feasibility of the method to enrich one leaflet with PIP_2_ and fluorescent TF-PIP_2_. We also quantitatively characterized the fluidity and asymmetry of the SLBs. We demonstrated the ability to fine-tune the amount of the asymmetrically incorporated lipids by incubating the SLB with varying concentrations of TF-PIP_2_. Our results suggest the potential application of this method to incorporate other lipids with sufficiently high CMC values, such as the other members of the phosphoinositide family and fluorescently labelled lipids^69,70^. While our study focused on the incorporation of lipids into SLBs, the possibility of adapting the method to other model membranes such as giant unilamellar vesicles (GUVs), large unilamellar vesicles (LUVs), and suspended planar lipid bilayers, or to the plasma membrane of cells, could open the potential applications to studies involving asymmetric membranes.

### Supporting information

Additional experimental data plots (PDF).

## Acknowledgment

We thank Frederic Pincet (ENS, Paris) for his help with the FRAP analysis and careful reading of the manuscript. We thank Patricia Bassereau and Julien Pernier for fruitful discussions. J.H, N.R., S.C. and F.-C.T. is a member of the CNRS consortium Approches Quantitatives du Vivant (AQV), Labex Cell(n)Scale (<u>ANR-11-889 LABX0038</u>) and Paris Sciences et Lettres (<u>ANR-10-IDEX-0001-02</u>). G.G. and F.-C.T. are funded by ANR grant <u>ANR-20-CE11-0010-01</u> (ActinFission). G.G. and F.-C.T., J.H., S.C., N.R., and J.B. are funded by a PSL-Qlife grant (MyoMemActin). JB is funded by grant R01GM115636 from NIH. T.L.A.N. is funded by École Doctorale Physique en Île-de-France (EDFPI) PhD fellowship. The authors greatly acknowledge the Cell and Tissue Imaging (PICT-IBiSA), Institut Curie, member of the French National Research Infractucture France-BioImaging (ANR10-INBS-04).

## Notes

### Competing Interest Statement

The authors have declared no competing interest.

### Summary of Updates

Figure 1 has new data: measurements of the CMC of PI(4,5)P2. New figure 2 combines the previous Fig. 2 and 4, and grouped data based on the duration between the SLBs samples prepared and the FRAP measurements. New Figure 4 combines the previous Fig. 6 and 7.

