## Supplementary material for "Asymmetric phosphoinositide lipid bilayers generated by spontaneous lipid insertion": SI

Supplementary information

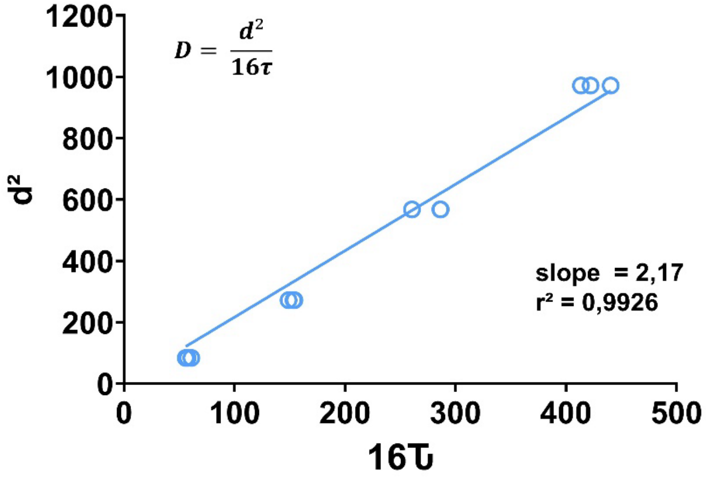


**Supplementary Figure 1.** Example of data fitting to determine the diffusion coefficient of lipids for one set of FRAP experiment. Each point represents a single FRAP experiment at a given area diameter d (µm) and characteristic time τ (sec) of a FRAP recovery. n = 12 FRAP measurements.

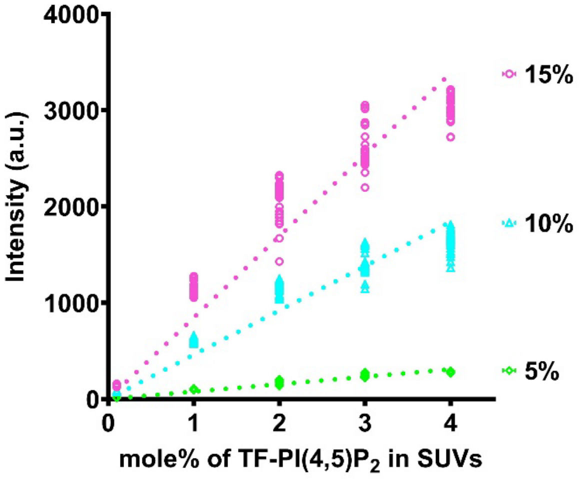


**Supplementary Figure 2. TF-PI(4,5)$P_2$ fluorescence intensities in sym-SLBs**. Sym-SLBs were generated by using SUVs containing concentrations of TF-PI(4,5)$P_2$ ranging from 0.1 mol% to 4 mol%. The SLBs were imaged using three different laser powers, 5, 10 and 15% by a 488 nm laser with a maximum laser power of 100 mW in a spin-disk confocal microscope. Dotted lines are linear fits to obtain $m_{sym-SLB}$ (intensity units per mol%, see Materials and Methods for details).

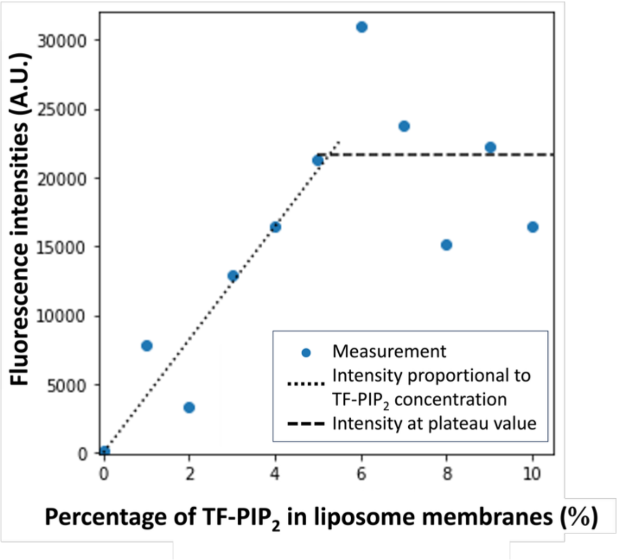


**Supplementary Figure 3. Fluorescence intensities of TF-PI(4,5)$P_2$ in POPC liposome membranes.** TF-PI(4,5)$P_2$ fluorescence intensity increases proportionally to its percentage in liposome membranes up to 5% TF-PI(4,5)$P_2$. Blue dots are measurements. Dotted line is the linear fit of the measurements for TF-PI(4,5)$P_2$ concentrations lower than 5%. Dashed line represents the plateau value of the intensities at TF-PI(4,5)$P_2$ concentrations higher than 5%.
